# Mental Chronometry in the Pocket? Timing Accuracy of Web Applications on Touchscreen and Keyboard Devices

**DOI:** 10.1101/639351

**Authors:** Thomas Pronk, Reinout W. Wiers, Bert Molenkamp, Jaap Murre

**Author notes:** Contact information for the corresponding author Name: Thomas Pronk, P.O. Box: 15969, 1001 NL, Amsterdam, The Netherlands, E-mail address.

## Abstract

Web applications can implement procedures for studying the speed of mental processes (mental chronometry). As web applications, these procedures can be administered via web-browsers on most commodity desktops, laptops, smartphones, and tablets. This approach to conducting mental chronometry offers various opportunities, such as increased scale, ease of data collection, and access to specific samples. However, validity and reliability may be threatened due to web applications on commodity devices having less accurate timing than specialized software and hardware. We have examined how accurately web applications time stimuli and register response times on commodity touchscreen and keyboard devices running a range of popular web-browsers. Additionally, we have explored the accuracy of a range of technical innovations for timing stimuli, presenting stimuli, and estimating stimulus duration. Results offer some guidelines as to what kind of methods may be most accurate, and what kind of mental chronometry paradigms may suitably be administered via web applications. In controlled circumstances, as can be realized in a lab setting, very accurate stimulus timing and moderately accurate Reaction Time (RT) measurements could be achieved on both touchscreen and keyboard devices. In uncontrolled circumstances, as may be encountered online, short stimulus durations (of up to 100 ms) may be inaccurate, and RT measurement may be affected by the occurrence of bi-modally distributed RT overestimations.

## Introduction

During the past decade, touchscreen devices (i.e., smartphones and tablets) have surpassed keyboard devices (i.e., desktops and laptops) in being the most frequently used devices for browsing on the internet (StatCounter, 2016). Web browsers offer platforms, based on widely supported open standards (such as HTML, CSS, and JavaScript) for deploying web applications to both touchscreen and keyboard devices. Hence, a research paradigm implemented as a web application can be deployed in the lab as well as on any commodity device, while also having the benefits of being based on open and durable standards. It has become more common to employ web applications for questionnaire research, but less so for mental chronometry (i.e. the study of the speed of mental processes). This is an important limitation, because psychological research increasingly employs mental chronometry to indirectly assess psychological constructs, which has been argued to help the validity of the assessments by reducing the influence of socially desirable answering (De Houwer, Teige-Mocigemba, Spruyt, & Moors, 2009; Greenwald, Poehlman, Uhlmann, & Banaji, 2009). If mental chronometry can reliably be conducted on touchscreen devices, this offers the opportunity to conduct such research on a wider range of samples than used to be feasible, such as inhabitants of emerging economies (Pew Research Center, 2016), and in a wider range of contexts, such as naturalistic settings (Torous, Friedman, & Keshavan, 2014).

One reason that mental chronometry on commodity devices via web applications has been limited, is doubt whether commodity devices have sufficiently accurate timing capabilities (Plant & Quinlan, 2013; van Steenbergen & Bocanegra, 2016). A range of studies have assessed the timing accuracy of web applications, but, to the best of our knowledge, only with keyboard devices. We make a general assessment of the technical capabilities of keyboard and touchscreen devices for mental chronometry paradigms in which a single static stimulus is presented to which a single response is registered. Such paradigms may require that stimuli are accurately presented for a specified duration and response times (RTs) are accurately measured. Factors that determine what level of accuracy is achieved, include the capabilities of the device, operating system, and web browser. Factors that determine what level of accuracy is required, include the demands of the particular paradigm under consideration, the degree to which systematic differences in accuracy across devices can be confounding variables, and to what extent these can be compensated by increasing the number of trials.

In two experiments, we examined the accuracy of stimulus presentation (Experiment 1) and the accuracy of RT measurement (Experiment 2) via external chronometry (i.e. measuring stimulus presentation via a brightness sensor and generating responses via a solenoid). Accuracy was examined on ten combinations of devices and browsers, formed by two touchscreen and two keyboard devices, each running a different operating system, and two or three browsers per device. Technical capabilities were evaluated for two research settings; a lab setting, in which device and browser could be controlled, and a web setting, in which they could not. For the former, the most accurate devices and browsers combination are evaluated, while for the latter, we evaluated variation across devices and browsers. As each experiment was designed and conducted in parallel, they are introduced jointly below.

With regard to stimulus presentation, web applications may occasionally realize shorter or longer durations than were requested (Barnhoorn, Haasnoot, Bocanegra, & Steenbergen, 2015; Garaizar & Reips, 2018; Garaizar, Vadillo, & López-de-Ipiña, 2014; Reimers & Stewart, 2015; Schmidt, 2001). Computer screens refresh with a constant frequency and presentation durations are typically counted in frames. The most common refresh rate is 60 Hz so that a frame then lasts about 16.67 milliseconds (ms). Timing errors can be expressed as *missed frames*, which is the number of frames realized minus the number of frames requested (Garaizar, Vadillo, & López-de-Ipiña, 2014; Garaizar, Vadillo, López-De-Ipiña, & Matute, 2014). We presuppose that a missed frame is problematic for mental chronometry paradigms in which stimuli are presented very briefly (e.g. 16.67 versus 33.33 ms) or for very specific durations, such as Posner tasks (Posner, 1980), stop signal tasks (Logan, Cowan, & Davis, 1984), and tasks using very briefly presented masked stimuli (Marcel, 1983). For longer durations, such as 250 ms, missed frames may be less problematic, as long as the realized duration does not differ too greatly from requested duration (e.g. 266.67 ms may be an acceptable deviation, but 350 ms may not).

Exploratively, we have evaluated a set of recently introduced methods for optimizing timing accuracy, sparked by developments in browser technology. We selected three methods with which a stimulus can be presented, based on manipulating the (1) opacity or (2) background-color Cascading Style Sheet (CSS) properties of a Hypertext Markup Language (HTML) element (Garaizar & Reips, 2018), and (3) drawing to a canvas element (Garaizar, Vadillo, & López-de-Ipiña, 2014). With regard to timing the onset and offset of stimuli, we selected two methods: (1) timing via requestAnimationFrame (rAF) (Barnhoorn et al., 2015; Garaizar & Reips, 2018; Garaizar, Vadillo, & López-de-Ipiña, 2014) for all three presentation methods and timing via CSS animation of the opacity and (2) background-position presentation methods (Garaizar & Reips, 2018). Additionally, we examined the accuracy of a range of measures for stimulus duration and RT via internal chronometry (i.e. measuring stimulus presentation and response registration using only the means available to the web application) in order to evaluate methods for improving timing accuracy or assessing accuracy afterward (Anwyl-Irvine, Massonnié, Flitton, Kirkham, & Evershed, 2018; Barnhoorn et al., 2015; Garaizar & Reips, 2018).

In Experiment 1, the accuracy of each of these presentation and timing methods were first assessed based on the proportion of cases in which the realized stimulus duration was exactly the number of frames as requested. Next, the most accurate method was further examined in terms of the distribution of missed frames, reporting on different methods if they produced notably different patterns of results. We compared the accuracy with which stimuli were presented for both brief and long durations — across devices and browsers — in order to assess to what degree mental chronometry paradigms may be affected that require very brief or specific stimulus durations.

With regard to RT measurement, research has indicated a noisy overestimation of RTs, with the mean and variance of overestimations varying across devices and browsers (Neath, Earle, Hallett, & Surprenant, 2011; Reimers & Stewart, 2015). In simulation studies, non-systematic overestimation of RT has generally been modeled as uniform distributions ranging up to 18 ms (Damian, 2010), 70 ms (Reimers & Stewart, 2015), and 90 ms (Brand & Bradley, 2012; Vadillo & Garaizar, 2016). Such RT overestimation was generally found to have a modest impact on a range of parameter estimation methods and designs, as most mental chronometry tasks measure within-subject differences in RT. However, systematic differences in RT overestimation between devices may form a confound when device preference systematically varies with a trait under study measured via absolute RT (but not RT difference) (Reimers & Stewart, 2015). Devices may also quantize RT into supra-millisecond resolutions (Reimers & Stewart, 2015). Even in this case, simulations revealed that resolutions of up to 32 ms may have little impact on the reliability of measuring RT differences (Ulricht & Giray, 1989).

Experiment 2 assessed how accurately web applications can measure RTs across devices and browsers. Both the average degree and range of RT overestimations were expected to vary substantially. In order to assess how well RT overestimations represented simulation assumptions, we examined distributions across devices, paying particular attention to the presence of any quantization. Similar to Experiment 1, only the most accurate method of timing and presenting stimuli was considered for further investigation; we will report on different methods if they produced notably different patterns of results.

Summarizing, the current study describes how accurate web applications on keyboard and touchscreen devices can present stimuli and register responses. For stimulus presentation, we examined the presence and magnitude of timing errors in terms of missed frames. For RT measurement we examined the accuracy with which RTs are measured, as well as distribution and quantization of RT overestimations. Exploratively, we examined a set of methods for improving timing accuracy based on different approaches to timing stimuli, presenting them, and measuring internal chronometry, so to assess the most accurate method that modern devices and browsers could offer.

## Methods

### Devices

Table 1 lists the characteristics of the devices used in the study. We selected one laptop for each of two popular keyboard operating systems (MacOS and Windows) and one smartphone for each of two popular touchscreen operating systems (Android and iOS). In the sequel, these devices will be referred to via their operating system. All four devices were normally in use as commodity devices by colleagues of the first author. We selected web browsers that were widely supported for each device: Chrome 71.0.3578.99 and Firefox 64.0.2/14.0 for all operating systems, and Safari 12 for MacOS and iOS. These browsers were selected for being relatively popular (StatCounter, 2018), still being actively developed at the time of the study, and each being based on a different browser engine, namely Blink, Gecko, and WebKit, respectively. Table 1 contains an overview of the specifications of each device. A device variable not included in our experiments was device load, as this seems to have only minor effects on modern devices (Barnhoorn et al., 2015; Pinet et al., 2016) and is difficult to manipulate systematically (and reproducibly) on touchscreen operating systems.

**Table 1.** Model, model number, and OS of each device.

| Device | Model | Model Number | OS |
| --- | --- | --- | --- |
| MacOS | MacBook Pro | A1398, EMC 2910 | MacOS 10.13.2 |
| Windows | ASUS Laptop | R301LA-FN218T | Windows 10.0.17134.523 |
| Android | Samsung Galaxy S7 | SM-G935F | Android 8.0.0 |
| iOS | iPhone 6S | MKU62ZD/A | iOS 12.1.2 |

### Design

During the stimulus presentation experiment, stimuli were presented for intervals of 16.67, 50, 100, 250, and 500 ms. As each device had a refresh rate of 60 Hz, these intervals corresponded to 1, 3, 6, 15, and 30 frames. The stimulus was a white square on a black background. The presentation experiment consisted of 120 sequences, in which each of the five combinations of timing and presentation methods was first prepared by adding the relevant HTML element to the web page, followed by one trial in each of the five intervals, followed by removing the HTML element. Each sequence was administered in one unique order out of the 120 possible orders in which five intervals could be arranged. Each sequence and interval was demarcated by 400 ms of background, followed by 400 ms of an inter-trial stimulus presented via a separate HTML element, followed by 400 ms of background. As the inter-trial stimulus was reliably presented for at least 200 ms across all devices, it could be used to match individual trials of each sequence on internal and external measures. During the reaction time experiment, a stimulus was presented until a response had been registered. For each of the five presentation methods, a sequence of 300 RT intervals was generated, consisting of each whole number of ms in the range 150 to 449 in a pseudo-random order. Each interval was distinguished by 1200 ms of background and each sequence was demarcated by 5000 ms of background.

### Measures

#### Stimulus presentation and timing

Three presentation methods were compared, in which stimulus onset and offset were realized by (1) manipulation of the opacity CSS attribute of a DIV element by changing it from 0 to 1 or from 1 to 0; (2) manipulation of the background-position of a DIV element by having a picture shift position such that a white or black part is presented; or (3) drawing a white or black rectangle on a Canvas element. Two timing methods were compared, in which stimulus onset and offset were: (1) timed via CSS animations that manipulated the appropriate CSS properties, or (2) timed by having a call from rAF manipulate the appropriate CSS properties or draw to a Canvas element, after a fixed number of frames. All manipulations were programmed via JavaScript, using the jQuery 3.31 library. The HTML, CSS, and JavaScript were configured according to best practices recommend in prior timing research (Garaizar & Reips, 2018): pictures were preloaded, each HTML element was laid out using absolute positioning in a separate layer, and CSS properties used for presenting stimuli were marked with the will-change property.

#### Internal chronometry

In Experiment 1, for all stimulus timing methods, four internal estimates of stimulus duration were compared. These measures, further described in Appendix A, were based on *rAF before created, rAF before called, raf after created*, and *raf after called*. For timing via CSS Animations, two additional estimates of stimulus duration were added, based on *animationstart/animationend created* and *animationstart/animationend called*. In Experiment 2, RT was internally measured as the time passed between stimulus onset and a keyboard or touchscreen response. For the moment of stimulus onset, the most accurate measure as found in Experiment 1 was selected. For the moment of response, two measures were compared, based on when a KeyboardEvent and TouchEvent were created or called.

#### External chronometry

Stimuli were detected via an optical sensor aimed at the top right of the device screen, which sampled luminance with a frequency of 3000 Hz. Stimulus onset and offset were defined as brightness increasing above and decreasing below 50% of maximum screen brightness, respectively. In Experiment 1, stimulus duration was measured as the number of frames that passed between stimulus onset and offset. In Experiment 2, an Arduino Leonardo microcontroller received a signal on stimulus onset, waited the number of ms as specified by each RT interval, and then sent a signal to trigger a solenoid. The solenoid was aimed at a touch-sensitive HTML element positioned at the bottom-right of the screen of touchscreen devices and at the Q key of keyboard devices. After triggering, the solenoid took 11 ms to go down and trigger a response on the key and touchscreen, as was measured with a second optical sensor at the bottom of the solenoid. Each interval between detecting stimulus onset and triggering solenoid was reduced by 11 ms to correct for solenoid delay.

#### Procedure

For each combination of device, browser, and experiment, the device was first prepared by installing updates to the operating systems and web browsers. Screen brightness was set to maximum, and screen-savers and notifications were disabled. Next, for each browser, the browser and experiment software was loaded, a minute waited (to allow background processes to reach a stationary level), and the experiment was started. Per combination of device and browser, Experiment 1 took about 80 minutes and Experiment 2 took about 40 minutes.

## Results

### Experiment 1

#### Accuracy per timing and presentation method

Table 2 shows the percentage of cases in which realized duration was exactly as requested (i.e. missed frames being 0) for each combination of device, browser, timing, and presentation method. Across all devices and browsers, except Mac OS Chrome, timing via rAF was more accurate than timing via CSS regardless of presentation method. Collapsed across all devices and browsers, timing via rAF and presenting via opacity (81.0%) was more accurate than timing via rAF and presentation via background-position (76.3%) or canvas (75.5%). When timed via CSS, presentation via background-position (23.7%) was more accurate than presentation via opacity (17.6%). There were substantial differences in accuracy across devices and browsers, ranging from a maximum of 59.3% accuracy on MacOS Chrome to near-perfect accuracy on iOS Chrome, Firefox, and Safari, and Windows Chrome. Hence, precise timing could be achieved for particular combinations of devices, browsers, timing, and presentation methods, but not consistently across all devices.

**Table 2.**
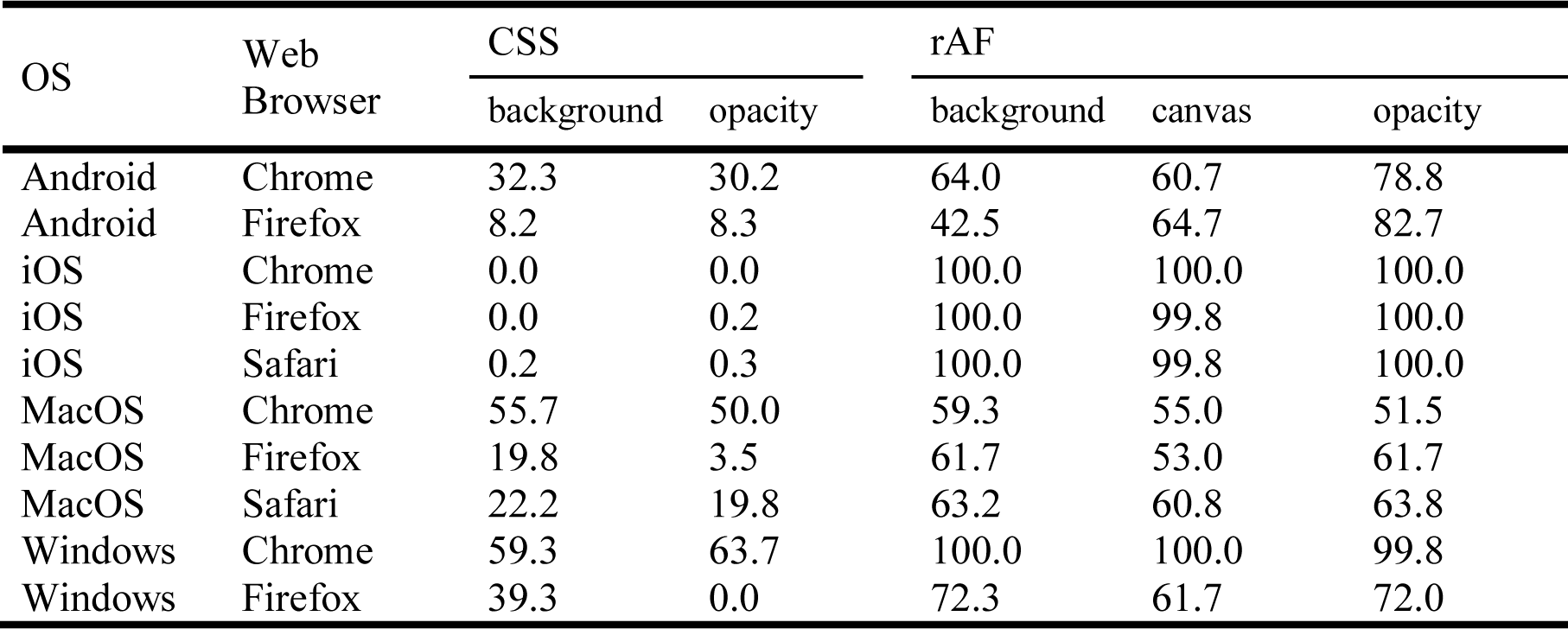
Percentage of trials in which realized duration was exactly as requested per device, browser, timing method, and presentation method

#### The accuracy of internal duration measures

We examined to what degree presentation duration could be estimated via internal chronometry, using four rAF-based measures and two animation-based measures. The distributions of rAF-based measures were strongly quantized with a resolution of 16.67 ms, while the distributions of the animation based measures were moderately so. In order to examine how accurately internal chronometry could predict realized duration, we quantized internally estimated durations into frames and calculated the percentage of trials in which realized duration was exactly the same as estimated via internal measures (i.e. missed frames being 0). For all devices, browsers, timing, and presentation methods, except MacOS Safari, rAF-based duration measures were more accurate than animation-based duration measures. Collapsed across devices and browsers, each of the four rAF-based measures was correct 75.8% to 76.3% of the trials, while animation-based measures were correct 34.8% to 34.9%. As rAF-based measures were more accurate than animation-based measures, these were selected for further analysis. Because each of the four rAF-based measures was similarly accurate, *rAF before created* was selected for further analysis. Table 3 shows the percentage of cases in which the missed frames were 0 for this measure per device, browser, timing method, and presentation method.

**Table 3.** Percentage of trials in which realized duration was exactly the duration as measured internally via rAF before created timestamps quantized into frames

| OS | Web Browser | CSS |  | rAF |  |  |
| --- | --- | --- | --- | --- | --- | --- |
|  |  | background | opacity | background | canvas | opacity |
| Android | Chrome | 58.3 | 70.7 | 64.0 | 60.7 | 78.8 |
| Android | Firefox | 53.5 | 80.7 | 45.5 | 64.5 | 82.5 |
| iOS | Chrome | 93.2 | 97.5 | 100.0 | 100.0 | 100.0 |
| iOS | Firefox | 94.5 | 95.2 | 100.0 | 99.8 | 100.0 |
| iOS | Safari | 93.5 | 94.3 | 100.0 | 99.8 | 100.0 |
| MacOS | Chrome | 56.3 | 50.2 | 59.5 | 55.0 | 51.5 |
| MacOS | Firefox | 60.7 | 64.8 | 61.7 | 53.0 | 61.7 |
| MacOS | Safari | 30.5 | 24.2 | 63.2 | 60.8 | 63.8 |
| Windows | Chrome | 99.8 | 99.5 | 100.0 | 100.0 | 99.8 |
| Windows | Firefox | 60.3 | 84.0 | 72.3 | 61.7 | 72.0 |

When timing via rAF, estimating stimulus durations via internal chronometry was similarly accurate as waiting a fixed number of frames was as at achieving stimulus durations. When timing via CSS, estimating stimulus duration via internal chronometry was more accurate than CSS animations were at achieving stimulus durations, but only for combinations of devices and browsers where timing via rAF already achieved near-perfect accuracy.

#### The magnitude of missed frames

The patterns of distributions of missed frames were consistent across presentation methods within rAF and animation-based timing methods but showed pronounced differences between timing methods. Therefore, below we report on the most accurate presentation method for timing via rAF (opacity) and for timing via CSS (background-color). Figure 1 shows the distribution of frame differences across devices, browsers, timing methods, and presentation intervals. There was variation in the sizes and signs of missed frames across devices, browsers, presentation methods, and intervals. Both iOS and Windows realized almost all stimulus durations within one frame of requested duration, as were stimuli timed via rAF on Android and MacOS Firefox. Timing via CSS yielded realized durations that were almost all consistently one frame longer than expected on iOS, as was the case on MacOS Firefox and Windows Firefox for one, three, and six frame intervals. Finally, note that stimuli requested for three or six-frame intervals were frequently presented too briefly on MacOS Chrome and MacOS safari. In fact, the majority of three-frame intervals were presented for only a single frame.

**Figure 1.**
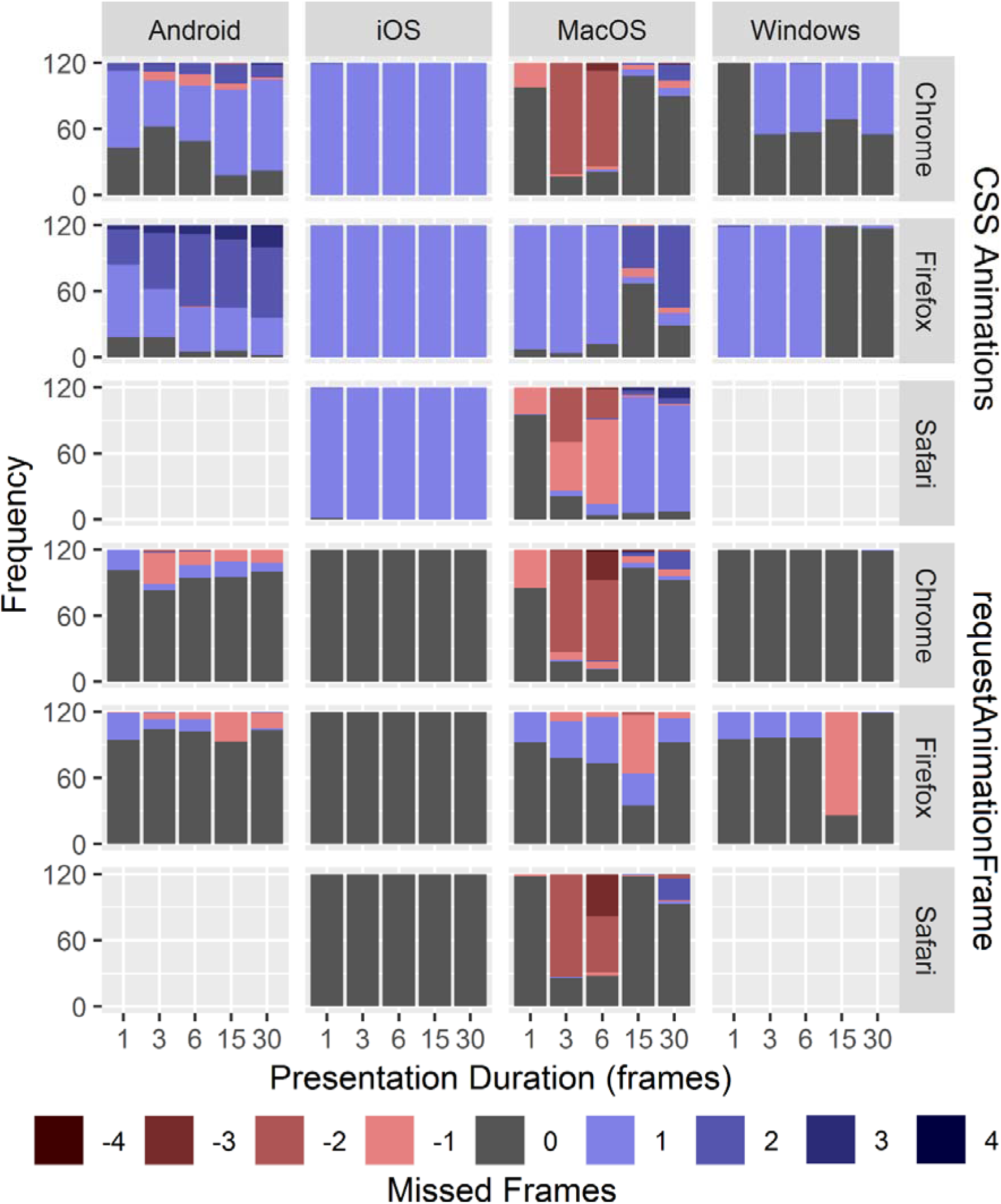
Stacked bar-charts of the frequency with which missed frames ranged -4 to 4 occurred per device, browser, and duration interval, for presentation via background-position and timing via CSS animations, and presentation via opacity with timing via rAF.

### Experiment 2

#### Absolute RT measurement

For RT measurements, only trials presenting stimuli via the most accurate timing method (rAF) and presentation method (opacity) were analyzed. As timestamp for stimulus onset, the internal measure *rAF before created* was used, and for response, *event created*, though the pattern of results was similar for other timestamps (all data and analysis scripts are available via the accompanying Open Science Foundation [OSF] website). RT overestimation was calculated as the difference between measured and realized RT. Table 4 shows the descriptives of RT overestimation per device and browser. Note that there were substantial differences between devices: mean RT overestimation on Android, iOS, and Windows ranged from 57 to 70 ms, with MacOS having higher overestimations, particular in combination with the Safari browser. The standard deviation (SD) of RT overestimation ranged from 5.7 to 8.1 across devices and browsers, except for Mac OS Firefox, which was particularly high, and Windows Chrome, which was particularly low.

**Table 4.**
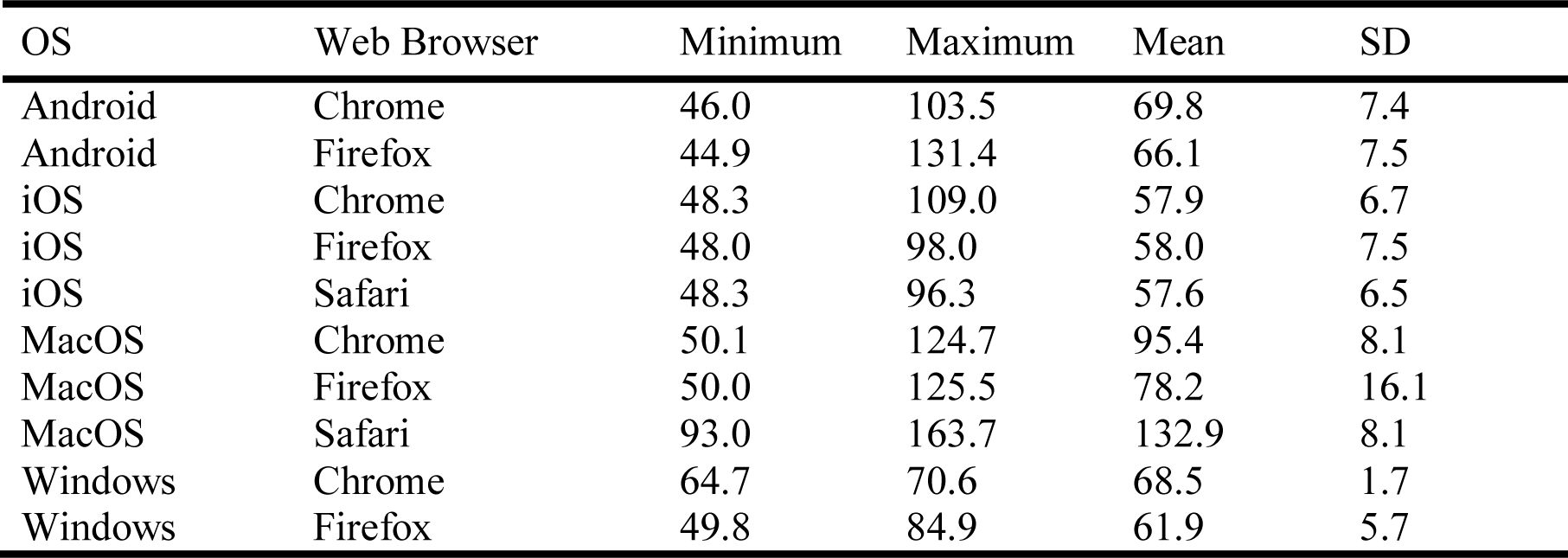
Descriptives of RT overestimations in ms per device and browser for stimuli that were timed via rAF and presented via opacity

#### Distribution and quantization of RT measurement

Inspection of scatterplots of measured RT versus external RT revealed quantization on iOS with a resolution of 60 Hz. Distributions of RT overestimations for most devices could be well described as having a normal to uniform distribution, with a small number of extreme outliers. However, Android showed some degree of bi-modality, while MacOS Chrome and MacOS Firefox showed pronounced bi-modal distributions (Figure 2). Fitting a mixture of two normal distributions via expectation maximization (Benaglia, Chauveau, Hunter, & Young, 2009) on RT overestimations of the most extreme case of bi-modality (MacOS Firefox) revealed two components, with mean (SD) values of 60.0 (12.4) and 89.1 (2.4) ms.

**Figure 2.**
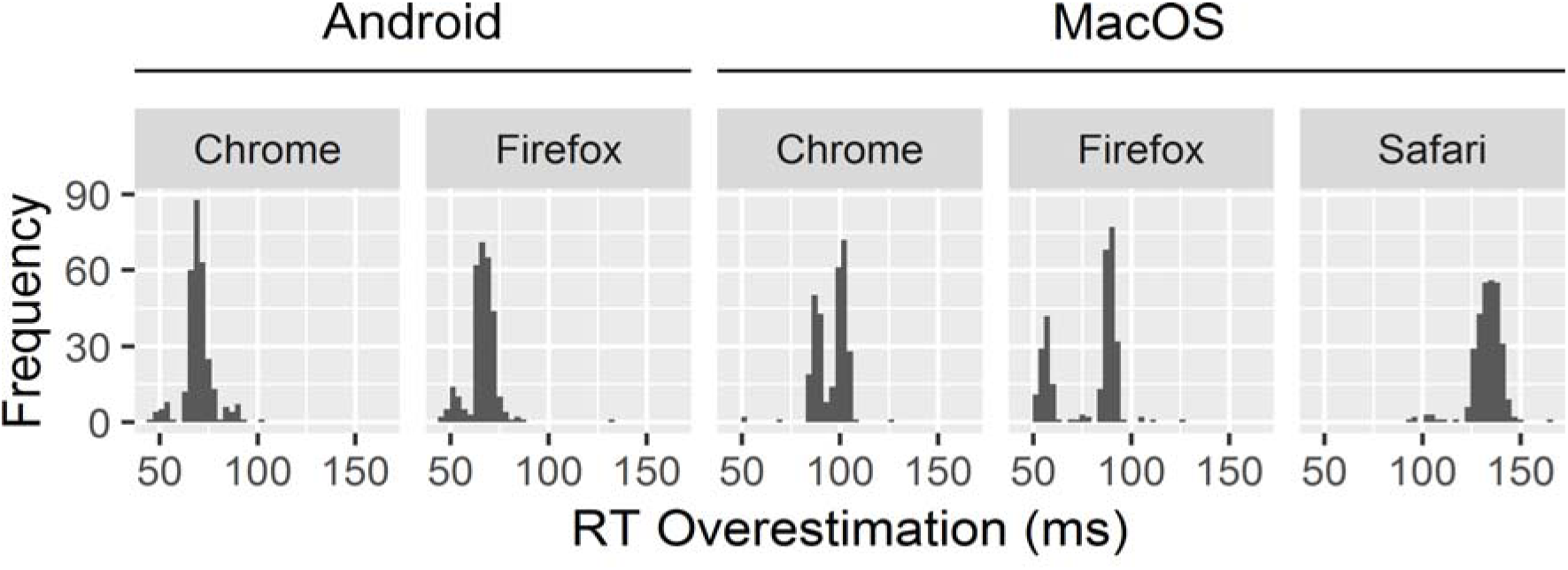
Distributions of RT overestimations on Android and MacOS.

## Discussion

We have examined how accurately web applications on touchscreen and keyboard devices present stimuli for a specified duration in Experiment 1 and measured RTs in Experiment 2. The results of each experiment are discussed below, followed by a general assessment of the technical capabilities of web applications for mental chronometry.

With regard to stimulus presentation, we first compare the results for different methods for timing and presenting stimuli, followed by an assessment of timing accuracy across devices and browsers. Timing via rAF was over 50 percentage points more accurate at realizing the exact stimulus duration that was requested than timing via CSS animations was. In part, this may be due to certain devices and browsers consistently presenting stimuli for one frame longer than requested. In those cases, requesting slightly shorter durations than was done in this study could improve the accuracy of stimuli timed with CSS animations to levels similar to rAF. However, such consistency was not found for all devices and browsers, so overall we recommend rAF for timing stimuli. We suspect that the inconsistencies in the behavior of CSS animations may be due standards for CSS animations still being a working draft (World Wide Web Consortium, 2018). When timing via rAF, presentation method had a relatively small effect on accuracy, with opacity outperforming background-position and canvas by up to five percentage points. As presentation methods were a relatively small factor in timing accuracy compared with timing methods, a researcher may consider choosing a presentation method based on practical considerations. For instance, canvas may be more suitable than opacity or background-position when dynamically generating stimuli. Also, note that a range of other presentation methods is supported by web browsers besides the three methods considered here, such as Scalable Vector Graphics and Web Graphics Library (Garaizar, Vadillo, & López-de-Ipiña, 2014). Future research could establish whether the findings reported here generalize to those presentation methods as well.

Internal chronometry measures of stimulus duration were similarly accurate in estimating realized stimulus duration as counting the number of frames was at achieving a requested stimulus duration. This finding is different from prior research (Barnhoorn et al., 2015; Garaizar & Reips, 2018), which may be due to differences in study aims and designs. The current study included a relatively large variety of devices and browsers and was the first to simultaneously compared timing stimuli by counting frames with estimating stimulus duration via internal measures. We found that for devices and browsers where stimulus timing by counting frames was near-perfect, internal measures for stimulus duration (e.g., JavaScript’s window.performance.now() high-resolution timer) were also near-perfect. Conversely, for devices and browsers where timing was less accurate, internal measures were less accurate as well. Hence, any increase in accuracy attributed to internal duration measures in previous studies may have been due to the devices and browsers on which the studies were conducted were already very accurate. While in the current study, internal chronometry could not provide any improvements in timing accuracy, internal chronometry may provide more general estimates of timing accuracy in a variety of other ways. For instance, an approach based on the regularity with which software events such as rAF occur (Eichstaedt, 2001) may be useful. Also, internal measures can be important for estimating the refresh rate of a device (Anwyl-Irvine et al., 2018). While beyond the scope of this paper, we hope to facilitate such approaches by making all data of the current study available for reanalysis; the URL to the Open Science Foundation containing all materials is listed at the end of this paper.

Based on the most accurate timing and presentation method found in this study (rAF and opacity), we assessed the accuracy with which keyboard and touchscreen devices can time stimuli. Some devices and browsers, both touchscreen and keyboard type, achieved near-perfect timing, namely iOS with Chrome, Firefox, and Safari, as well as Windows with Chrome. Hence, in settings where device and browser can be controlled, web-applications can be as accurate as specialized software. Most devices and browsers achieved most presentation durations within one frame of requested duration, though on a keyboard device, two browsers tended to present durations of up to six frames (100 ms) too briefly. Hence, when the device and browser cannot be controlled, the reliability and validity of mental chronometry paradigms that require brief presentations may be affected. An analysis with operating system and browser type is recommended in cases where absolute timing accuracy is is of high importance (e.g., [near] subliminal presentation).

With regard to the accuracy of RT measurements, different internal measures for RT showed a similar pattern of results. Quantization of RT into 60 Hz was found on one device, which may be acceptable, at least for RT differences (Ulricht & Giray, 1989). RT overestimation varied across devices, similar to what was found in previous research (Neath et al., 2011; Reimers & Stewart, 2015). On three devices, two touchscreens and one keyboard, the shape of RT distributions had a smaller range and was more normal than the uniform distributions assumed in simulation studies (Brand & Bradley, 2012; Damian, 2010; Reimers & Stewart, 2015; Vadillo & Garaizar, 2016), but with a comparatively small range. Hence, when the device that is administering mental chronometry can be controlled, RT may be measured quite accurately, but not at the level that specialized hardware and software, such as button boxes under Linux, can provide (Stewart, 2006). On particular combinations of devices and browsers, namely MacOS with Chrome and Firefox, RT overestimations were bi-modally distributed with centers that could differ up to 30 ms. While in this case, the range of device noise was still relatively small compared to simulation studies, an occasional occurrence of bi-modal noise may affect parameter estimations in unexpected ways. Again, we recommend an analysis with operating system and browser type as mentioned above for the case of brief stimulus presentation. Except for the cases where RT overestimations were bi-modally distributed, the number of trials commonly used in mental chronometry tasks (of 32 and more) is sufficient for device noise to negligibly affect statistical power.

In summary, in controlled settings, web applications may time stimuli quite accurately and register reaction times sufficiently accurately when a constant overestimation of RTs is acceptable. In uncontrolled settings, web applications may time stimuli insufficiently accurately for mental chronometry paradigms that require brief stimulus presentations. Also, the occurrence of bi-modal noise, which was found on iOS browsers, may affect parameter estimates in some experimental paradigms.

Web applications offer a means to deploy studies inside and outside of the lab. Frameworks are being developed that make it increasingly easier for researchers to deploy mental chronometry paradigms as web applications (Anwyl-Irvine et al., 2018; De Leeuw, 2015; Henninger, Mertens, Shevchenko, & Hilbig, 2017; Murre, 2016). Studies of timing accuracy suggest limits to what may be achieved but also introduce a range of technical innovations for achieving higher accuracy. The experiments reported in this paper examine a range of these technical innovations in order to offer some guidelines on optimal methods. A sample of ten combinations of devices and browsers was studied, so that these guidelines and what level of accuracy can be achieved may be generalized with some confidence.

The results in this study may be representative for web browsers, as the three browsers selected in this study represent a large majority of browsers used online (StatCounter, 2018). The sample of four devices, though selected for being in use as commodity devices and each having a different operating system, was nevertheless small compared to the variety of devices available to web applications. However, if the systematic differences in accuracy across devices and browsers that was found in this study would replicate in a larger sample, then online studies could correct for it by detecting participants’ device and browser.

Overall, touchscreen devices seem technically capable of administering a substantial number of mental chronometry paradigms, when taking some limitations and best practices into account. As smartphone ownership and internet connectivity are becoming ubiquitous, this offers various opportunities for administering mental chronometry on large scales and outside of the lab. By implementing reaction time tasks as web applications, they are based on durable and open standards, allowing a single implementation to be deployed on desktops, laptops, smartphones, and tablets. We hope that this paper helps to answer doubts about the timing accuracy of such an approach.

## Supporting information

Appendix A. An Overview of Timestamps Provided by Web-browsers that can be used for Internal Chronometry

## Acknowledgments

We thank Jeanine Baartmans, Marilisa Boffo, Bruno Boutin, and Jasper Wijnen, for making their personal devices available for this study.

## Open Practices Statement

Study documentation, scripts of procedures of web applications and solenoid timing, data, and analysis scripts, are available at https://osf.io/4fq3s.

