## Appendix A. An Overview of Timestamps Provided by Web-browsers that can be used for Internal Chronometry for "Mental Chronometry in the Pocket? Timing Accuracy of Web Applications on Touchscreen and Keyboard Devices"

Thomas Pronk, M Sc[1]

Reinout Wiers, Prof Dr[1]

Bert Molenkamp, B Eng[1]

Jaap Murre, Prof Dr[1]

**Affiliations**

[1] University of Amsterdam, Faculty of Social and Behavioural Sciences Amsterdam, Department of Psychology, Amsterdam, The Netherlands

**Contact information for the corresponding author**

Name: Thomas Pronk

P.O. Box: 15969, 1001 NL, Amsterdam, The Netherlands

### Introduction

JavaScript web-applications have an event-driven architecture (Mozilla, 2019a), which means that the application is structured as reactions to events created by the web browser. These events occur, for instance, when the device screen is refreshed or when a participant undertakes an action such as tapping a touchscreen or pressing a key. An application can react to an event by registering one or more functions as event handlers, which are then called when an event of a particular type is created. Via the web-browser, timestamps can be obtained that indicate when an event happened. These timestamps could be referred to as internal chronometry as they measure time using only the means available to the web application, as opposed to external chronometry, which measures time using external sensors. Internal chronometry has been used to estimate response times (RTs) (Barnhoorn, Haasnoot, Bocanegra, & Steenbergen, 2015; Neath, Earle, Hallett, & Surprenant, 2011; Pinet et al., 2016; Reimers & Stewart, 2015), but more recently, also to estimate and improve the accuracy with which stimuli are presented for a specified duration (Anwyl-Irvine, Massonnié, Flitton, Kirkham, & Evershed, 2018; Barnhoorn et al., 2015; Garaizar & Reips, 2018). This appendix contains an overview of timestamps that may be used for internal chronometry and a timeline illustrating when each timestamp is logged.

Traditionally, timestamps that were used to establish the moment of stimulus onset, stimulus offset, or a response, were recorded at the moment an event handler was *called* (Barnhoorn et al., 2015; Neath et al., 2011; Reimers & Stewart, 2015). More recently, the moment an event was *created* (or started firing) has been proposed as a more accurate alternative (Anwyl-Irvine et al., 2018; Garaizar & Reips, 2018). The rationale for the latter is that the timestamp recorded at event creation is less subject to delays than the timestamp recorded at event handling. We have extended this rationale to the timing of requestAnimationFrame (rAF) relative to screen refreshes, as well as CSS animation and response-related events. Listing 1 contains JavaScript that illustrates how to obtain the *created* and *called* timestamp in relation to rAF and to Events, with the following sections explaining each in more detail.

*Listing 1. JavaScript code that illustrates how to collect created and called timestamps for a rAF handler and event handler*

```
// raf_called and raf_created
requestAnimationFrame(function (raf_created) {
     raf_called = window.performance.now();
});
// event_called and event_created
document.addEventListener(“touchstart”, function (event) {
     event_called = window.performance.now();
     event_created = event.timeStamp;
});
```

### Timestamps related to requestAnimationFrame

Figure 1 shows a timeline of relevant timestamps. Four of these are related to a technology called requestAnimationFrame (rAF) (Mozilla, 2019b), which will be explained first. Computer screens refresh at a constant rate. When a web application draws graphics, they are not immediately displayed but will be at the next refresh. Via rAF, a web-application can synchronize with the refresh such that sufficient time is available for drawing all required graphics before the next refresh takes place. While the web application cannot, at present, obtain a timestamp of the moment that a refresh takes place, it can obtain timestamps in relation to rAF. Drawing and displaying graphics via rAF at refresh X then consists of the following steps, with related timestamps in italics:

1. After refresh X - 1, rAF starts firing (*rAF before created*)
2. An event handler provided by the application is called (*rAF before called*) that draws the graphics
3. The graphics are displayed at refresh X.
4. After refresh X, rAF starts firing again (*rAF after created*)
5. An event handler provided by the application may be called again (*rAF after created)*

When a device is performing optimally, one could expect that rAF starts firing briefly after each refresh and that the function provided by the application is called briefly after rAF starts firing. Hence, the timestamp that rAF starts firing after refresh X (rAF after created) may, in optimal circumstances, be closest to the moment the refresh takes place.

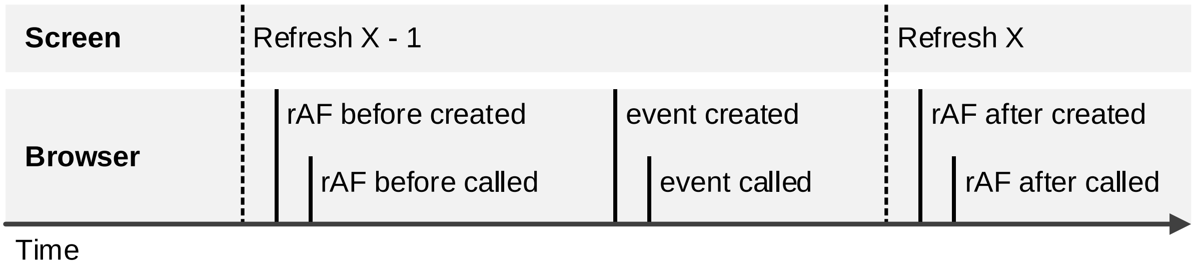


*Figure 1. A timeline of timestamps related to an event happening at refresh X or between refresh X - 1 and refresh X*

#### Timestamps related to Animation Events

In addition to the rAF-related timestamps, Figure 1 shows two *event* timestamps. These timestamps concern events that may take place between *rAF before called* and *rAF after called*. When drawing graphics via CSS animations, an animationstart and animationend event can be used to inform the web application when a CSS animation started and finished. We note that the sequence of timestamps of *animationstart* and *animationend* events relative to rAF may vary across web-browsers. For example, on Android Chrome 71.0.3578.99 we observed that *animationstart* and *animationend* tended to be between *rAF created* and *rAF called*. However, on MacOS Safari 12 we observed that timestamps for *animationstart* took place briefly after *rAF before called*, while timestamps for *animationend* took place briefly before *rAF after created*.

### Estimating Stimulus Duration via Timestamps

In sub-optimal circumstances, one of the steps required for displaying graphics may be delayed. When drawing graphics that represent stimulus onset or offset, this may result in a stimulus being displayed for too short for too long. A possible method for counteracting this problem proceeds as follows: at the rAF event handler that draws stimulus onset, a timestamp is recorded. Each time rAF calls the event handler, the number ms passed since the timestamp of stimulus onset is calculated. If the number of ms exceeds the requested stimulus duration minus 5 ms, graphics are drawn for stimulus offset. This way, a delay in stimulus onset can be adjusted for by a delay in stimulus offset. As timestamp for stimulus onset and offset, Barnhoorn and colleagues (2015) used *rAF before called*, while Garaizar and Reips (2018) and Anwyl-Irvine and colleagues (2018) used *rAF before created*. The validity of these methods hinges on the assumption that timestamps collected during stimulus onset and offset accurately estimate the actual duration that the stimulus was displayed. In total, four estimates of stimulus duration can be considered, based on the time passed between stimulus onset and offset as measured via *rAF before created*, *rAF before called*, *rAF after created, rAF after called.* Timestamps related to *rAF before* may be more useful than timestamps related to *rAF after,* as the former can be used to correct for delays in stimulus onset, while the latter cannot. However, timestamps related to *rAF after* may be more accurate, as they may be recorded closer to the refreshes corresponding to stimulus onset and offset than timestamps related to *rAF before* are. When presenting stimuli via CSS animations, two additional methods for estimating stimulus duration are available, based on *animationstart/animationend created* and *animationstart/animationend called.*

### Estimating RT via Timestamps

When a participant responds by pressing a key, a KeyboardEvent of type *keydown* is created, and when a participant responds by pressing a touchscreen, a TouchEvent of type *touchstart* is created. By comparing timestamps recorded at stimulus onset with timestamps recorded at *keydown* or *touchstart* events, RT can be estimated. Similar to rAF, timestamps at *keydown created* or *touchstart created* may be closer to the moment the response took place than the timestamp at *keydown called* or *touchstart called*, and so be a more accurate estimate of RT. Regardless, since timestamps recorded at stimulus onset may not reflect the precise moment a screen is refreshed and a stimulus is presented, some noise is inherent. Similarly, different devices and web-browsers may introduce a range of delays between the moment a response is given and when it is registered (Neath et al., 2011; Reimers & Stewart, 2015).
